# Mass Mortality of Wood Frogs (*Rana sylvatica*) in the Ozark National Forest: Is it Acid Rain?

**DOI:** 10.1101/338970

**Authors:** Malcolm L. McCallum, Tracy Klotz, William Stephens, Jennifer Bouldin, Ken Gillespie, David Feldman, Benjamin A. Wheeler, Stanley E. Trauth

## Abstract

We describe observations of a continuing mass mortality event primarily involving wood frogs, but involving other amphibians to a lesser degree. The investigation took place from Spring 2000 through Spring 2004. No definitive correlations between environmental variables and mortality could be identified. Forensic analysis could not isolate causal pathogens. Although mortality fluctuated during the study, it may have spread to other species. Our report identifies population level problems in the eastern part of the Ozark National Forest but is unable to identify a cause. Future studies that more thoroughly address contaminants, pathogens/parasites, and other potential environmental problems in the Ozark National Forest are warranted.

## Introduction

The wood frog (*Rana sylvatica*) is a widespread anuran, occurring as far north as the Arctic Circle and as far south as Alabama; however, most of its range is restricted to the northern reaches of North America. The populations located on the Ozark Plateau are non-continuous with the rest of the species’ range (Trauth et al. 1995; Trauth et al. 2004). Such disjunctions are important to conservation of a species because of their propensity to contain unusual phenotypes adapted to these refugia (Boughey, 1973; Southerland, 2000).

Wood frogs employ an explosive breeding strategy in which nearly the entire population breeds in the same night, usually after the first warm rain in January or February (Bellis, 1962). These frogs are listed as a species of concern by the Arkansas Natural Heritage despite their state conservation status (S4, common secure), and global rank (G4 globally secure, rare in parts of its range) partly because they are currently a state inventory element. Mysterious deaths of wood frogs were first documented during the Spring 2000 breeding chorus in Stout Pond. This pond is located in the Sylamore Ranger District of the Ozark-St. Francis National Forest. Mass mortality events such as this die-off are identified as one of the primary observations related to amphibian populations declines (Reaser, 1996) associated with current biodiversity losses in amphibians and other groups (McCallum 2007; McCallum 2015).

This report outlines the results obtained from surveying populations across the Sylamore District, and from monitoring Stout Pond for four years. It also provides recommendations regarding what steps are required to identify the cause/s, and to manage these important forest resources.

## Materials and Methods

The methodologies we employed to investigate wood frog die-offs were divided into several areas: (1) surveys for dead and dying frogs, (2) basic water quality measurements, (3) acute and chronic toxicity bioassays.

We traveled to the Sylamore Ranger District of the Ozark National Forest every rainy night in the spring that could be characterized as a typical stimulus of wood frog breeding. On nights when wood frog breeding occurred, we surveyed ponds for dead frogs. All ponds were searched thoroughly by walking transects across the entire pond basin and surrounding bank area. Dead frogs were collected and either fixed in 10% formalin and preserved in 70% ethanol, or packed in ice for transport back to the ASU Museum of Zoology herpetological collection where they were held at −70 C according to the United State Geological Survey’s (USGS) Amphibian Research and Monitoring Initiative (ARMI) Standard Operating Procedures (SOPs). Frozen and preserved frogs were sent to the USGS Wildlife Health Center (Madison, Wisconsin) for forensic pathology.

Stout Pond was monitored by Trauth since 1985. McCallum & Trauth continued monitoring from 2000-2003. Each year we recorded the species that were observed and noted if they were dead and alive. Representative voucher specimens were collected and then deposited in the ASUMZ herpetological collection each year. In 2001 and 2002 approximately 15 students accompanied us during the chorus. In 2003, Lisa Irwin (USFWS Conway Office, AR), Kelly Irwin (AGFC state herpetologist), and four students assisted during the wood frog chorus.

Water samples and seven wood frog egg masses were collected from Stout Pond and a single reference pond (near Blanchard’s Springs Caverns) in 2002, placed on ice and transported to the Arkansas State University Ecotoxicology Facility and Museum of Zoology where basic water chemistry was employed. These samples were used in acute and chronic toxicity bioassays (APHA, 1998; USEPA, 1993). The tests used were the *Pimephales promelas* 48 hr acute toxicity bioassay and the *Chironimus tentans* 10-day growth and survivorship bioassay. Four egg masses (3,472 eggs) were placed together in 6 L of water from stout pond. Three clutches (936 eggs) were placed together in 2 L of dechlorinated tap water. Hatching rate in each water sample was recorded.

Water Quality measurements were made at 12 ponds in the vicinity of Stout Pond near Fifty Six (Stone County, Arkansas) in 2003. Measurements included pH, DO, water temperature, air temperature, and conductivity. Observations of live and dead amphibians, amphibian eggs, evidence of potential predators, and freezing conditions were recorded. Water samples and three amplectant pair of wood frogs were collected and returned to the ASUMZ herpetological collection. These breeding pairs were allowed to oviposit in dechlorinated tap water. Twenty eggs randomly chosen from among all three amplectant pairs were placed in three replicates of six different hatch test regimes. All treatments were housed in 10 × 15 × 30 cm plastic aquaria containing 2000 mL of water. Where sediment was included in the treatment, 500 g was employed. Treatment 1 contained water and sediment collected from Stout Pond. Treatment 2 contained sediment from Stout Pond with tap water. Treatment 3 contained only water from Pavilion Pond (ASU campus). Treatment 4 contained only water from Stout Pond. Treatment 5 contained only unbuffered tap water. Treatment 6 contained tap water and Pavilion Pond sediment. Eggs were allowed to develop and biomass production, hatching success, and survivorship were recorded and compared between groups.

## Results

### Environmental Data and Mortality

Wood frogs bred on 26 February 2000, 9 February 2001, 30 January 2002, and 20 February 2003. On all visits ringed salamander (*Ambystoma annulatum*) adults and larvae were alive and evident. On 26 February 2000 we observed 140 dead wood frogs at Stout Pond. Of these, 53.6% (n = 75) were males and 46.4% (n = 65) were females. Among the females, 89.2% (n = 58) were gravid and 6% (n = 4) were found dead in amplexus with live males and 4.6% (n = 3) were dead with extra-peritoneal egg extrusions into the thigh and loin regions. The average male SVL was 51.2 mm (range = 45.9 −58.2 mm) and average female SVL was 60.1 mm (range = 54.7-66.0 mm). A single portion of a ringed salamander was found dead in the communal egg mass.

On 9 February 2001 Stout Pond was experiencing extreme drought conditions and only about 10-15% of its basin contained water. The main breeding chorus occurred around 2 am according to nearby residents; we arrived later that evening at 6 pm. The previous 24 hours were unseasonably warm (near 60 F). We observed 27 dead males and 8 dead females. Some males amplexed dead females, and some females were amplexed with dead males. We also observed amplectant pairs with both individuals dead. The temperature began to drop after we arrived and was below freezing by 9 pm. The communal egg mass filled the entire water filled area (approximately 5 m × 6 m) of Stout pond.

On 30 January 2002 we arrived at Stout Pond at 5:41 pm. The temperature = 3 C, wind speed = 0.9 m/s. The water had a pH = 5.09, DO = 6.78 mg/L, and conductivity = 0.016 μS. We observed 35 dead frogs and a communal egg mass approximately 20 m × 10 m. The pond was completely filled with water.

It harbored at least 100 spotted salamander (*Ambystoma maculatum*) egg masses, each containing white embryos, apparently dead.

In 2003 we visited Stout Pond again, this time there were 262 dead wood frogs and we observed two dead adult spotted salamanders. Frogs were observed dying in the chorus by Kelly (Arkansas Game and Fish) and Lisa Irwin (Arkansas Department of Environmental Quality) prior to my arrival. Conductivity, pH, and observations of live/dead amphibians are documented for Stout Pond and 11 other ponds in the Sylamore Ranger District. A view of one area taken through a single camera shot to show the extent of mortality at Stout Pond is shown in Figure 1 (also see Fig. 2). There were also dead frogs with signs of red leg (Fig. 3)

**Figure 1.**
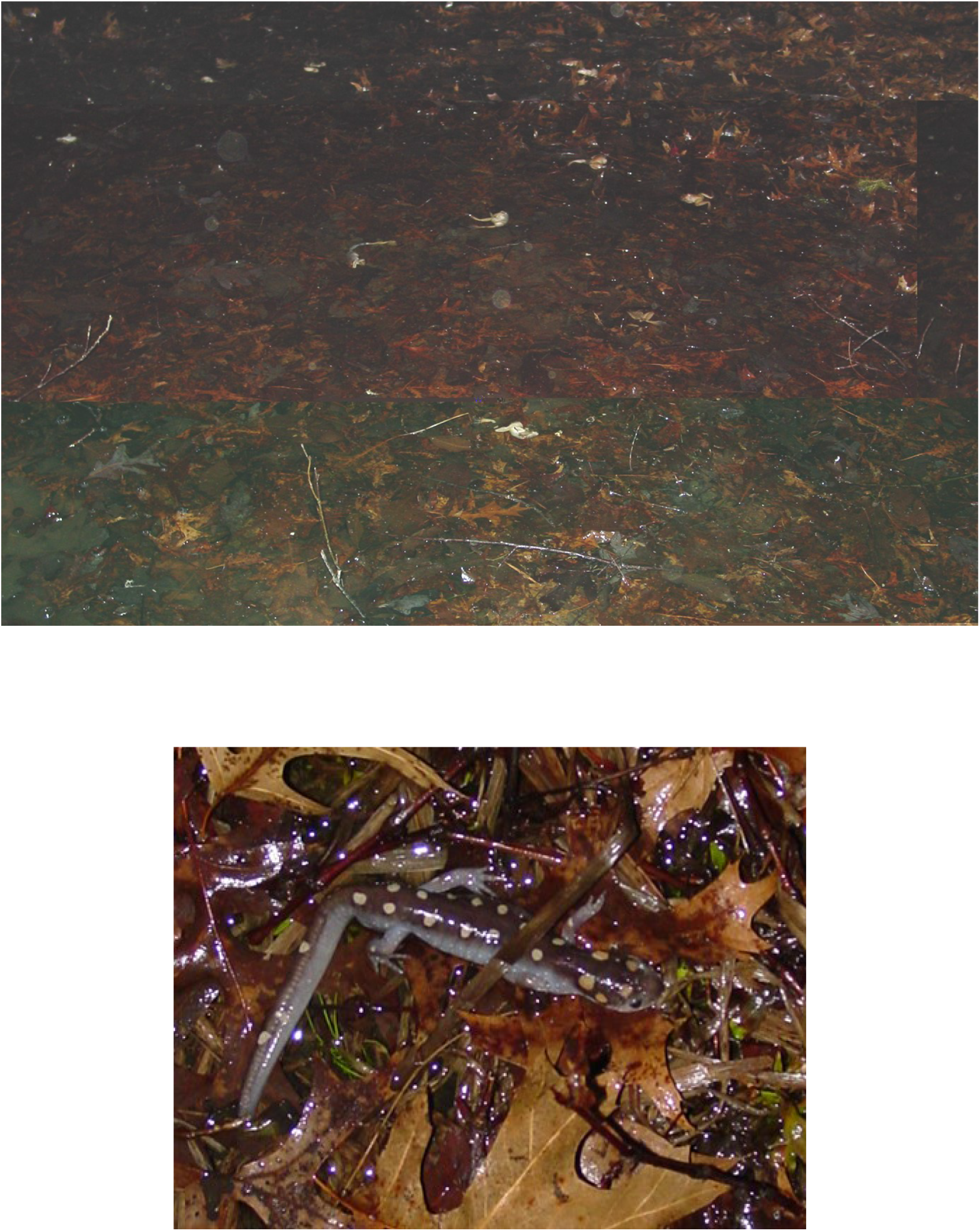
Photograph of dead Wood Frogs (*Rana sylvatica*) and a dead Spotted Salamander (*Ambystoma maculatum*) on the southeast shore of Stout Pond, Ozark National Forest, Arkansas (Syalmore District).

**Figure 2.**
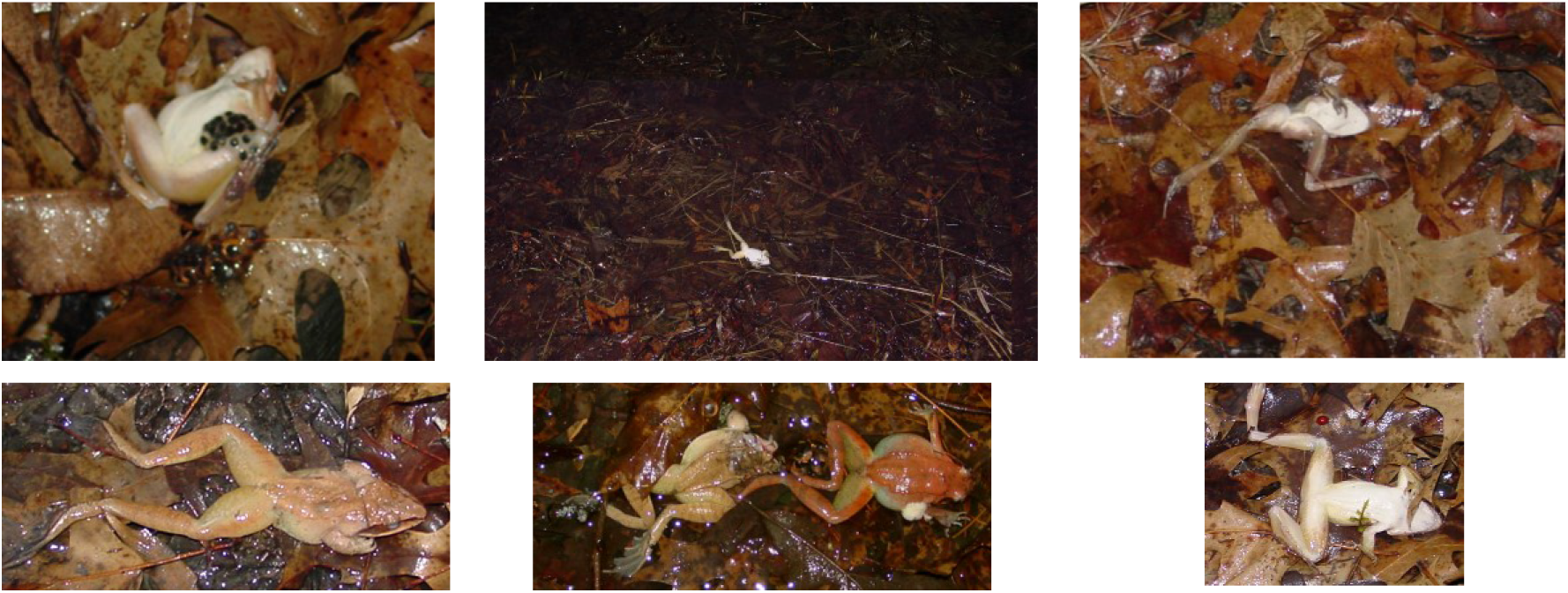
Disposition of representative Wood Frogs (*Rana sylvatica*) from mass mortality event in the Sylamore District of the Ozark National Forest, Arkansas.

**Figure 3.**
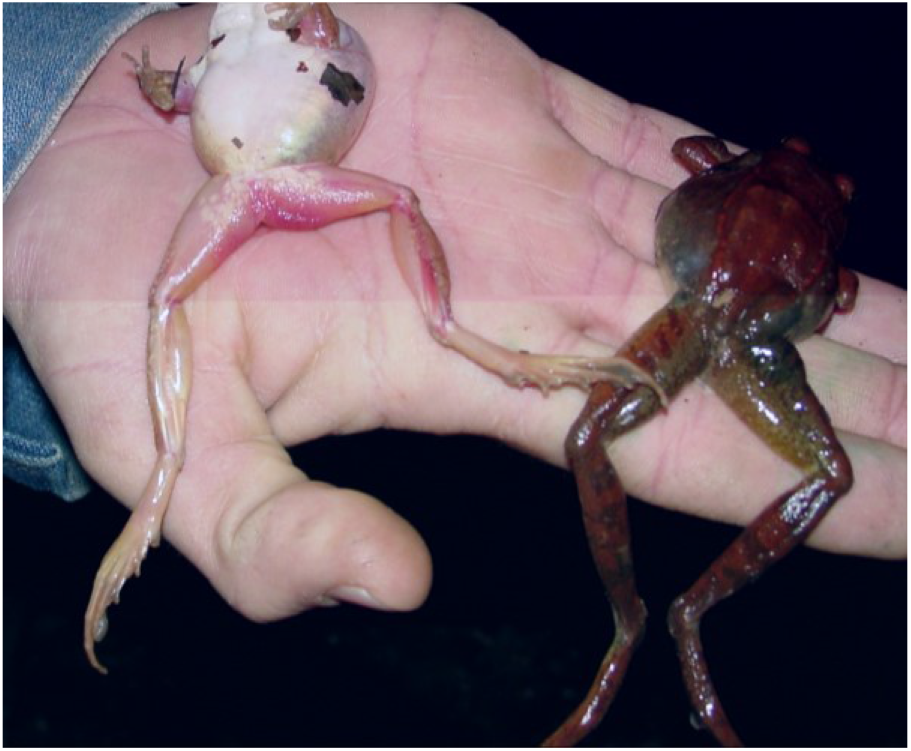
Dead wood frogs with signs of Red Leg Disease, note red flush in inguinal region and apparent bubbles under the skin on specimen on the right.

Dead frogs were handled according to ARMI SOP’s and mailed to David Green at the USGS Wildlife Health Center in Madison, WI. They screened the specimens for known pathogens. Mechanical trauma was observed in 4/12 specimens. Cestodes, trematodes, and Myxozoans were observed in 3/12 individuals. All 12 specimens observed showed severe atrophy of fat bodies.

In 2003 we searched for dead amphibians at 12 additional ponds in the Sylamore and conducted water quality measurements on these ponds. Five of these ponds contained dead eggs and/or amphibians and one had no signs of amphibian activity. The results of these searches are given in Table 1. No significant relationship between pH and signs of dead amphibians (G = 0.783, df = 1, *P* = 0.0.376) when all species were lumped together. Egg mortality was marginally related to pH (G = 1.548, df = 1, *P* = 0.213) and more strongly to conductivity (G = 2.978, df = 1, *P* = 0.084). The occurrence of dead amphibians as a group was closely associated with low conductivity (G = 3.611, df = 1, P = 0.057). Our data suggest that there may be some weak interaction between pH and conductivity (G = 4.339, df = 2, *P* = 0.114) affecting the presence of dead amphibians and there eggs. *Pimephales promelas* 48 hour acute toxicity showed no toxicity to either Stout Pond water or a single unaffected reference pond where living wood frogs were observed. The *Chironimus tentans* 10 day growth and survival test showed a borderline level of impairment for the sediment (P = 0.043) and no impairment for water.

**TABLE 1.** Water chemistry and Mortality at 12 ponds in the Sylamore District of the Ozark National Forest. Key: W = wood frog, S = spotted salamander, P = Spring Peeper (*Pseudacris crucifer*), SE = spotted salamander egg mass, WE = wood frog egg masses, B = bullfrog, SL = ringed salamander larvae, BL = bullfrog (*Rana catesbeiana*) larvae.

| Pond # | pH | Conductivity | Species Present | Mortality/Comments |
| --- | --- | --- | --- | --- |
| **Stout** | 5 | 28.5 | W, S, SE, WE | 262 W, WE, 2 S |
| **1** | 5.78 | 30.5 | W, P | 4 W, 1 P |
| **2** | 6.07 | 78.5 | few eggs (no ID) |  |
| **3** | 5.6 | 21.2 | SE | SE WE (most fungal) |
| **4** | 4.9 | 17.1 |  |  |
| **5** | 5.16 | 14.1 | W, WE, SL | WE frozen |
| **6** | 4.48 | 17.7 | WE, SE | 1/3 WE, SE |
| **7** | 6.09 | 120.4 | SL | Bare earth eroding into pond |
| **8** | 5 | 16.2 | B | 1 juvenile B, 1 adult B |
| **9** | 7.8 | 74.8 | BL |  |
| **10** | 4.5 | 34.3 | WE (fresh laid) |  |
| **11** | 5.1 | 25.5 | 1 live P |  |

### Laboratory Hatch Tests

Eggs from the 2002 hatch test showed only 8.7% (302/3472) egg survivorship in water from Stout Pond; whereas, there was 70.1% (656/936) egg survivorship in dechlorinated tap water. In 2003 the more refined hatch test showed differences in overall survivorship through time (*F* = 24.10, df = 10, *P* < 0.001) and among treatments (*F* = 31.35, df = 5, *P* < 0.001). There was an interaction between these treatments was of marginal importance (*F* = 1.27, df = 50, *P* = 0.143). Stout Pond sediment with Stout Pond water with stout pond sediment had 40.9 % survivorship (mean = 8.18 tadpoles). Stout Pond sediment with tap water had 45% survivorship (mean = 8.79 tadpoles) and Stout Pond water without sediment had 40 % survivorship (mean = 8 tadpoles). Water from Pavilion Pond showed 78.8 % survivorship (mean = 15.76 tadpoles).

Stout Pond water and sediment showed significantly higher biomass production than the other treatments (F = 19.90, df = 8, *P* < 0.001). Stout Pond water and sediment produced a biomass mean = 3.63 g (SD = 0.22). Stout pond sediment with tap water produced tadpole mean biomass = 2.06 g (SD = 0.45). Stout Pond water without sediment produced a tadpole mean biomass = 2.01 g (SD = 0.22). Pavilion Pond water produced a tadpole mean biomass = 2.26 (SD = 0.07). There were no differences among treatments for individual tadpole mass (F = 0.38, *P* = 0.918).

## Discussion

Our observations suggest amphibian mortality in the Ozarks is not simply a sporadic event located at a single pond, but rather a more widespread phenomena occurring throughout at least the Sylamore District, if not all of the eastern Ozarks. The Ozarks are a biodiversity hot bed, and numerous endemic amphibians reside there. If a pathogen or environmental problem is accentuating the levels of amphibian mortality in this region, many of these species previously thought to be secure could be approaching levels of concern. The mortality in this study does not appear related to a pathogen or chemical contaminant. The observance of individuals with symptoms of red leg is implicative and previous die-offs of wood frogs in other regions have involved red leg pathogens (Nyman, 1986). However, no pathogens were identified in our frogs. Further, the apparent damage observed to many dead amphibians (Trauth et al. 200 was traced primarily to feeding by snails scavenging the corpses. Additionally, the past drought conditions are not likely to play much of a role in egg mortality in our settings (Forrester et al., 1988).

Regular observations of amphibian mortality from 2000 – 2004 at Stout Pond may stem from acid rain deposition. The Ozark region has long received sulfur and nitrogen deposition originating from Louisiana and Texas, and it receives 14.1 – 17 lbs/acre of sulfur each year (Hunt and Heilman 1999). Even though the impacts from acid rain in the Ozarks have been lower than expected (Marion et al. 2014); the association of acidic conditions and amphibian mortality cannot be over-looked. Acid deposition was expected to drop with the 1990s amendments to the Clean Air Act, but the problem persists.

The relationship between wood frog egg mortality and conductivity has been previously reported (Gascon et al. 1987; Peirce et al. 1984). When soils buffering capacity is exhausted, runoff does not pick up pH elevating cations, thus rain-fed ponds develop low pH and conductivity values like those we observed. Other areas of the Ozark Plateau show that acid rain rapidly exhausts the buffering capacity of Ozark soil (Spratt 1997). The conductivity (28 – 120 μS) and pH (pH = 4.8 – 7.8) observed in the ponds we visited are vastly lower than observed in similar ponds (pH = 6.4 – 7.4; Conductivity = 75 – 253 μS) in the immediate vicinity decades earlier (Chapman et al. 1982). It is possible that the surface soils in this region are beginning to lose their buffering capacity contributing to egg deaths. In fact, the only pond with a pH above 6.09 was located immediately adjacent to a limestone outcrop. This calcium load undoubtedly helped maintain its higher pH. Regardless, the acid deposition alone may be sufficient to acidify these ponds.

Wood Frogs continue to hold on in the Ozark Plateau; however, one must question if the organism communities in this biodiverse region will increasingly feel the impacts of continued acid deposition. Until emissions from distant, out-of-state sources are reigned in, it is difficult to imagine that a viable solution for these problems will reveal itself.

## Acknowledgments

We would like to thank Dr. David Greene (USGS Wildlife Health Center) for conducting forensic analysis and Jerry L. Farris (Arkansas State University) for use of field and laboratory equipment. Additional thanks to the Sylamore District of the US Forest Service for financial assistance through a series of Challenge-Cost Shares.

